# Native spike flexibility revealed by BSL3 Cryo-ET of active SARS-CoV-2 virions

**DOI:** 10.1101/2023.10.10.561643

**Authors:** Hideo Fukuhara, Hisham M. Dokainish, Shunsuke Kita, Koshiro Tabata, Akira Takasu, Juha T. Huiskonen, Yuki Anraku, Tomo Nomai, Taiyo Someya, Toshiya Senda, David I. Stuart, Michihito Sasaki, Yasuko Orba, Yasuhiko Suzuki, Hirofumi Sawa, Katsumi Maenaka

**Affiliations:** Division of Pathogen Structure, International Institute for Zoonosis Control, Hokkaido University, Kita-20, Nishi-10, Kita-ku, Sapporo 001-0020, Japan; Laboratory of Biomolecular Science and Center for Research and Education on Drug Discovery, Faculty of Pharmaceutical Sciences, Hokkaido University, Kita-12, Nishi-6, Kita-ku, Sapporo 060-0812, Japan; Global Station for Biosurfaces and Drug Discovery, Hokkaido University, Kita-12, Nishi-6, Kita-ku, Sapporo 060-0812, Japan; Division of Molecular Pathobiology, International Institute for Zoonosis Control, Hokkaido University, Kita-20, Nishi-10, Kita-ku, Sapporo 001-0020, Japan; Institute for Vaccine Research and Development (HU-IVReD), Hokkaido University, Kita-21, Nishi-11, Kita-ku, Sapporo 001-0020, Japan; Structural Biology Research Center, Institute of Materials Structure Science, High Energy Accelerator Research Organization (KEK), Tokyo 135-0063, Japan; Institute of Biotechnology, Helsinki Institute of Life Science HiLIFE, University of Helsinki, P.O. Box 56, 00014 Helsinki, Finland; Department of Accelerator Science, School of High Energy Accelerator Science, SOKENDAI (the Graduate University for Advanced Studies), 1-1 Oho, Tsukuba, Ibaraki 305-0801, Japan; Faculty of Pure and Applied Sciences, University of Tsukuba, 1-1-1 Tennodai, Tsukuba, Ibaraki 305-8572, Japan; Division of Structural Biology, University of Oxford, The Wellcome Centre for Human Genetics, Headington, Oxford OX3 7BN, UK; Chinese Academy of Medical Sciences Oxford Institute, University of Oxford, Oxford OX3 7FZ, UK; Diamond Light Source Ltd, Harwell Science & Innovation Campus, Didcot OX11 0DE, UK; Nuffield Department of Medicine, Wellcome Centre for Human Genetics, University of Oxford, Oxford OX3 7BN, UK; Division of Bioresources, International Institute for Zoonosis Control, Hokkaido University, Kita-20, Nishi-10, Kita-ku, Sapporo 001-0020, Japan; One Health Research Center, Hokkaido University, Kita-19, Nishi-9, Kita-ku, Sapporo 001-0020, Japan; Global Virus Network, Baltimore, Maryland, USA; Faculty of Pharmaceutical Sciences, Kyushu University, Fukuoka, 812-8582, Japan; Research Center for Vaccine Development, National Institute of Infectious Diseases, Japan Institute for Health Security, Tokyo 162-8640, Japan

**Keywords:** SASR-CoV-2, Spike protein, Cryo-ET, Vaccine design, Conformational dynamics, Native active virions

## Abstract

Understanding the molecular properties of severe acute respiratory syndrome coronavirus 2 (SARS-CoV-2) is crucial for tackling future outbreaks. Current structural knowledge of the trimeric spike protein relies on truncated recombinant proteins and/or inactivated full-length forms, which may suffer from overstabilization. Here, we apply cryo-electron tomography (cryo-ET) at a Biosafety Level 3 facility to study the virus structure in its native, active state. The virus particles exhibit variable shapes and sizes with diffusible spikes, with the majority in typical prefusion conformations. Notably, we identified unprecedented, a transient open-trimer prefusion states, revealing a hidden flexibility with opened S1 conformation. Subtomogram averaging of the prefusion spikes indicates a loosely packed trimeric architecture that may facilitate the formation of open-trimer state. A cryo-EM map of recombinant Omicron BA.2.75 spike protein further confirms the presence of this loosely packed trimer as a minor conformational state. The observed dynamics uncover conserved cryptic regions that can be targeted for broadly effective vaccines. Structural analysis of active viruses profoundly impacts our understanding of the overlooked fusion mechanism and vaccine, antibody/drug design.

## Introduction

The recent SARS-CoV-2 pandemic has highlighted the profound impact of infectious diseases on people’s lives^1^. The emergence of new variants hinders the efficacy of developed vaccines and therapeutic antibodies, emphasizing the threat of future outbreaks^2,3^. The spike protein decorates the virus surface and plays central roles in viral entry and immune shielding mechanisms^4,5^. The spike protein is a trimeric glycoprotein, wherein each protomer is formed of two subunits (S1 and S2) with a transmembrane region^6^. Numerous studies elucidated spike protein structures and dynamics, paving the way for mRNA vaccine developments^6–10^. Specifically, cryogenic electron microscopy (cryo-EM) has been used to determine structures of the recombinant spike protein in wild-type and emerging variants^3,6–13^. Moreover, cryogenic electron tomography (cryo-ET) has enabled the study of full-length spikes in the inactivated viruses, unraveling the flexibility of its stalk region and the protein distribution on the virus surface^7,14–20^. Structural studies provided fundamental information on the virus molecular structure, architecture, interactions, and dynamics^4^.

Despite the enormous amount of structural information available on SARS-CoV-2 virus and its spike protein, our understanding of the virus in its active form is still lacking^21^. In fact, the majority of cryo-EM studies are performed using a truncated ectodomain of the spike protein, which includes several mutations (e.g., 2P and 6P proline mutations) designed to stabilize the prefusion state^22,23^. Accordingly, several approved vaccines include similar proline mutations, for improving production and stabilization^24^. Indeed, the proline mutations may induce local rigidity altering the nearby secondary structure^25^. Likewise, to reduce biological hazards, chemical fixation is usually applied in cryo-ET experiments^16,21^. Chemicals such as paraformaldehyde (PFA) are known to react with amino residues and may cause morphological changes that alter protein structures^21,26^. In addition, electron (E)-beam inactivation was recently shown to be damaging to spike protein structures^16^. According to previous structural studies, the spike protein has a reasonably stable prefusion state with a number of conformational states that are governed by the motion of its S1 receptor binding domains (RBDs) from Down to Up states^13^. In contrast, hydrogen deuterium exchange mass spectrometry (HDX-MS) study showed that the prefusion state of the spike protein is quite dynamic, highlighting the existence of a form of the prefusion state in which parts of S2 are exposed^25^. Likewise, previous high-speed atomic force microscopy (AFM) experiment indicated the lateral motion of RBD, suggesting higher level of flexibility than simple RBD Down to Up motion^27^. Furthermore, the binding of some antibodies to the S2 subunit in the prefusion state suggests the formation of an unknown open trimer structure^28^. Collectively, previous studies showed structural flexibility at several levels using artificially stabilized proteins and inactivated viruses. However, nature of such flexible conformations remain unknown as native images of spike proteins are still missing.

Understanding the differences between the current structural knowledge of SARS-CoV-2 that may suffer from overstabilization, and the active form is crucial for future vaccine and antibody/drug development. The rational of this work is to investigate the structure of SARS-CoV-2 virions in their active state with minimal experimental handling. The infectivity and pathogenesis of viruses depend on the interplay and physicochemical properties of the host and the virus. For viruses, such as SARS-CoV-2, there is a wealth of evidence that the glycoprotein responsible for the dominant neutralizing antibody responses and for cell attachment and fusion is a dynamic structure, whose dynamics modulate its function. To address this point, we set up the high-end cryo-electron microscope in a Biosafety level 3 (BSL3) laboratory to employ cryo-ET and subtomogram averaging (STA)^29^ to unravel the structure of the spike protein using active viruses. In this experiment, neither chemical fixation nor ultracentrifugation were applied, to avoid spike structure alteration or detachment. This study represents a structural analysis of SARS-CoV-2 in its active form. Hence, the current study shows that, while the vast amount of reported works is on non-physiological states, the implication of this study is important to understand the precise functional mechanism relevant for anti-drug and vaccine design.

## Results

### BSL3 High-end Cryo-electron Microscope

We set up the latest generation of cryo-electron microscope (300 kV), Krios G4 (Thermo Fisher Scientific) with a Falcon 4 direct electron detector and a Selectris energy filter in the management area for BSL3 pathogens in the International Institute for Zoonosis Control, Hokkaido University (Figure 1–figure supplement 1). The detail of BSL3 cryo-EM system and its biosafety setup are described in the following section, but briefly, virus vitrification using EM GP2 (Leica) and autogrid assembly were all performed in the safety cabinet. The autogrids were transferred to Krios G4 with originally designed cryotransfer system and cryo-ET data were collected (Figure 1–figure supplement 1). The medium containing the infectious SARS-CoV-2 (ancestral strain WK-521) was collected from VeroE6/TMPRSS2 cells 18 hours post-infection. It was subsequently employed for plunge freezing, followed by cryo-ET. Thus, the virus preparation did not include ultracentrifugation or chemical fixation process to avoid any kind of artificial effects. However, the sample still experienced external forces during the filtration and surface tension in the blotting processes. Furthermore, virions were produced in VeroE6/TMPRSS2 cells which overexpress TMPRSS2 to maintain ancestral SARS-CoV-2 features^44^.

**Figure 1.**
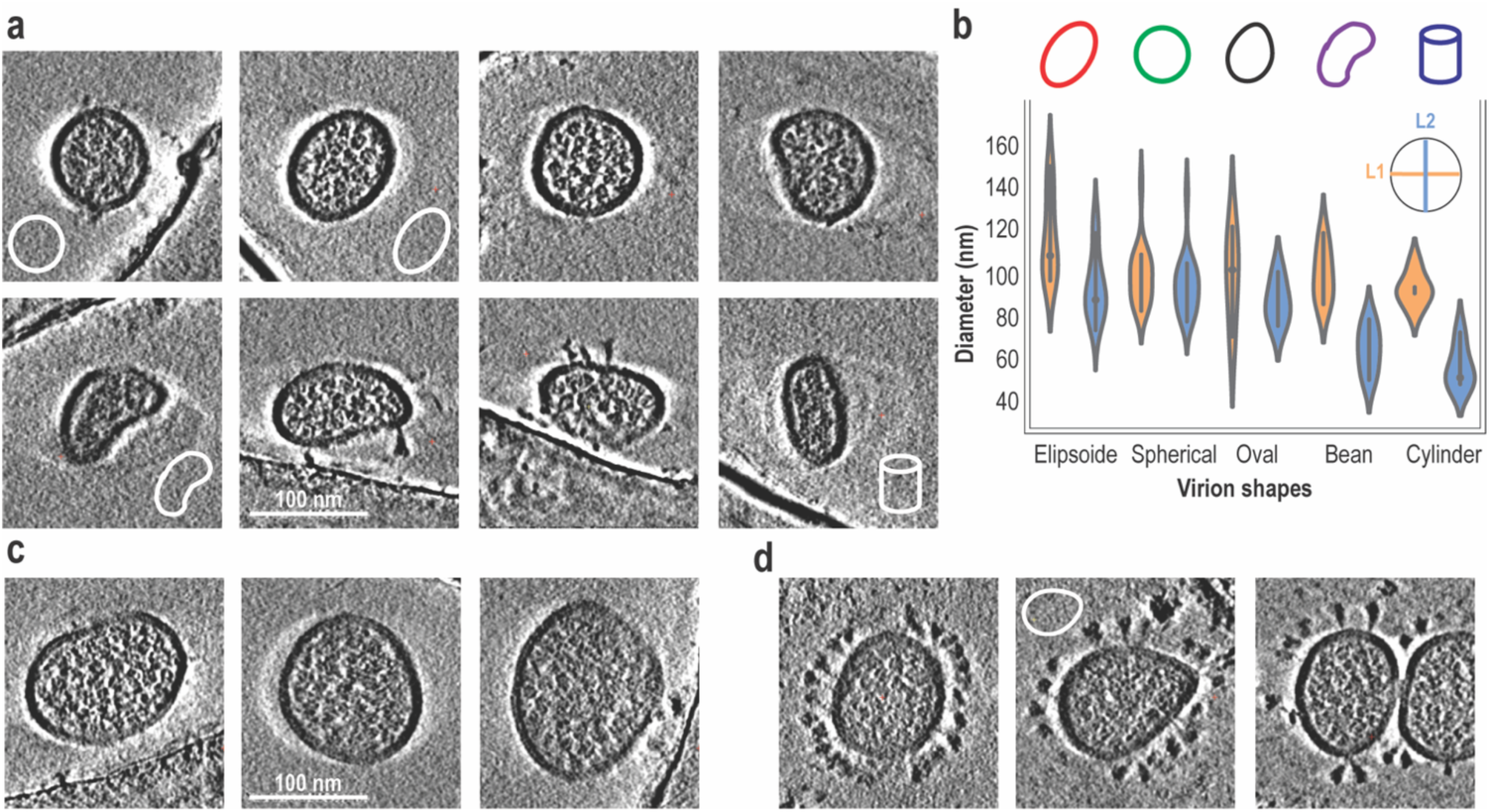
Pleomorphology of active virus particles. a) Central sections (7.5 nm thick) through the cryo-ET tomograms of the ancestral virions. b) Violin plot representation of the two longest diameters (L1 and L2) in 46 virus particles. c) Tomograms slices of three virions with very large diameters (Left: L1:139, L2:102; Middle: L1:125, L2:119; Right: L1:150, L2:125). d) Central slices of virions on carbon film. Scale bar of 100 nm is also shown. **Figure 1–figure supplement 1.** BSL3 Cryo-EM facility at Hokkaido University and Cryo-ET analysis scheme. **Figure 1–figure supplement 2.** Particle morphology and ice thickness. **Figure 1–figure supplement 3. Morphology of inactive virions.** **Figure 1—video 1.** 2D and 3D Reconstruct of several tomograms of active SARS-CoV-2 virions. **Figure 1—video 2.** Large variations in virion sizes. **Figure 1—video 3.** Two virion particles undergoing fusion. **Supplementary file 1A, Table 1.** All particles diameter measurements (L1 and L2) in nm. **Supplementary file 1B, Table 2.** Comparison with previous Cryo-ET studies of SARS-CoV-2. **Supplementary file 1C,Table 3**.Cryo-ET data acquisition and reconstruction statistics.

### Installation and handling of cryo-EM at BSL3 area

To achieve cryo-EM observation of native SARS-CoV-2, there were some challenges to overcome as follows: how to decontaminate the cryo-EM itself for maintenance, how to prevent viral leakage during vitrification, clipping, and cryo-transfer, and how to supply liquid nitrogen into the P3 area. To decontaminate the cryo-EM, we introduced a manufacturer-designed heating system inside the enclosure. Due to the prior consideration, SARS-CoV-2 delta variant was confirmed to be inactivated with heating at 60 °C for 120 min. The equipped heating system can maintain the inside of the cryo-EM enclosure at 60 °C for more than 3 h. All tubes connected to the vacuum pumps were equipped with inline HEPA filters. In addition, disinfection of the entire room is accomplished with vaporized hydrogen peroxide (VHP) fumigation (STERIS). To supply liquid nitrogen, two tanks were prepared inside and outside P3 area. To prevent accidental backflow, the tanks are normally disconnected. When the remaining amount of liquid nitrogen inside the tank becomes low, we connect these tanks manually and transfer the liquid nitrogen while monitoring the pressure to maintain one-way flow. Next, EM GP2 was used for grid vitrification. A large Class II safety cabinet with sufficient space for placing EM GP2 and clipping grids was custom-ordered. Ethane gas was supplied through a HEPA filter into a safety cabinet. The cryo-transfer system was modified based on the manufacturer’s NanoCab and connected via HEPA-filtered suction unit during transfer and docking so that airflow is not directed to the outside. This system was first checked for operation with a 3D-printed model, and the final parts were then replaced with PTFE plastic and stainless steel. After constructing the above system, we tested operations and its completeness using coronavirus HCoV-229E. Some concerned positions were wiped off during the sequential operation and checked for viral contamination by HCoV-229E, and no contamination was detected in the whole process without a positive control

### The Pleomorphic Virus Particles Morphology

Early cryo-ET study of the inactivated SARS-CoV-2 demonstrated the formation of a dominant ellipsoidal or a spherical-shaped viral envelope, with an average of 96.6 ± 11.8 nm for its longest axis^17^. In contrast, a more recent cryo-ET study suggested the formation of a flat cylindrical shape, in ancestral, alpha, beta and delta variants, with an average diameter of 102 nm^15^. Notably, a later cryo-ET study of the delta variant has shown the formation of fused virus particles, where one envelope encloses two sets of RNPs^16^. In summary, while the majority of cryo-ET studies of inactivated virion suggested spherical/ellipsoidal shape, few studies have suggested some level of pleomorphism^30^.

In this study, 42 tomograms of the active ancestral virus were recorded and reconstructed, including 46 virus particles (e.g., Figure 1—video 1). A tilt angular step of 1°, covering a range from −60° to +60° was used, resulting in 121 tilt images per tomogram. Figure 1 shows a variety of envelope shapes, including dominant spherical (L1: 99 ± 13.6 nm, L2: 94 ± 13.6 nm) and ellipsoidal shapes (L1: 117 ± 19.7 nm, L2: 94 ± 15.9 nm), as well as oval (L1: 99 ± 21.5 nm, L2: 87 ± 10.6 nm), cylindrical (L1: 93 ± 7.9 nm, L2: 57 ± 10.1 nm) and bean-like (L1: 100 ± 12.7 nm, L2: 65 ± 10.9 nm) envelops (summarized in Supplementary file 1A, Table 1). The measured diameters of the two longest axes (L1 and L2) indicate a large variation in the virus size, wherein the maximum axis (L1) can vary from 72 to 150 nm (Figure 1—video 2). Remarkably, very large virus particles with L1:150, L2:125 (Figure1c right) and L1:139, L2:102 nm (Figure1c left) were observed. Such large particles may indicate the fusion of two virus particles in ancestral strain, as observed in delta variant^16^. Figure 1–figure supplement 2a and Figure 1—video 3 also show the observation of two virus particles apparently undergoing a fusion process.

To unravel the nature of these large virions, we estimated the number of RNP from the central slice of very large virion in Figure1c (right), observing 44 RNP. This reflects the unusually large genome compared to typical size virion, which contain around 15 RNP along central slice and overall ∼45 per virion. Figure 1–figure supplement 2b shows that there is no correlation between the observed morphology and ice thickness, confirming the minimal effect on the virions’ shape. It also shows that the third axis is correlated with the particle size. Collectively, unlike previous results from inactivated viruses that exhibit a dominant morphology (Supplementary file 1B, Table 2), our results suggest a higher level of flexibility, adaptability, and pleomorphism on virus shape in response to surrounding environments, demonstrating the effect of inactivation on virion shapes and sizes.

To further elucidate the effect of inactivation on virion morphology, we collected a data set using viruses treated with the same protocol as the native virus preparation, except for inactivation using paraformaldehyde. In this dataset, 69 virions from 55 tomograms were constructed. Figure 1–figure supplement 3a confirms the effect of inactivation, wherein the virion diameters are less diverse, occurring at a lower range with an average of 91.8 ± 12.3 nm and 82.2 ± 11.4 nm for L1 and L2, respectively, compared to active virions (103.4 ± 16.9 nm and 85.8 ± 19.1 nm). Only two virions with relatively large diameters were observed, as shown in Figure 1–figure supplement 3b, though the longest observed diameter is much smaller than that of large active virions. Figure 1–figure supplement 3c also shows that two main shapes, either spherical or ellipsoidal, were observed. As pointed out in previous studies^21,26^, inactivation has a direct effect on virion shape and flexibility, where native form shows higher polymorphism.

### Spike Distribution and Arrangements

In the current dataset, one-third of the particles (14 out of 46) were observed on the carbon film. These particles tend to include a higher density of spike proteins, in comparison to the ones in the grid holes. Note that, in the absence of fiducial markers, we found that carbon regions facilitate more reliable patch tracking, which makes the alignment quality of particles on the carbon comparable to those in the ice. On the carbon, spike proteins are pushed to the edge of the flattened particle, reflecting high diffusibility within the native viral lipid membrane (Figure 1d). Such high diffusibility may allow the virus to relocate more spikes at the interface upon encountering natural host receptor, Angiotensin-converting enzyme 2 (ACE2). Previous experimental and coarse grained computational studies showed that spike trimers might move laterally^31,32^. On the other hand, within grid holes, spike proteins are more dispersed on the surface, where fewer spikes can be identified in most cross sections. In general, the number of spikes varied significantly among virus particles, where one virion within a hole contained 22 spikes, while others exhibited 14 or 15 spikes. The total number of spikes per virion is consistent with previous estimates from fixed cryo-ET data^17,30^.

Interestingly, Figure 2a shows that spike proteins can exist in close proximity to form clusters containing 2 or more spike proteins (dimer or trimer like) (Figure 1—video 1 right). Similar complexes were only observed in full-length spike proteins from the Novavax vaccine^33^. Furthermore, aside from the close proximity observed in the head region, analogous behavior is also evident in the stem region, as depicted in Figure 2–figure supplement 1a. Likewise, the postfusion state was found to occur in clusters (Figure2b) that may facilitate membrane fusion by attaching several fusion peptides (FP) at the same time. Comparison with previous studies also shows that the pre-/post-fusion ratio is highly affected by preparation and inactivation, while active virions show 85 to 15% ratio, β-Propiolactone (BPL) inactivation increases post-fusion to 48%, and paraformaldehyde drastically reduces post-fusion occurrence, as observed in this study as well as previous works (see Supplementary file 1B, Table 2).

**Figure 2.**
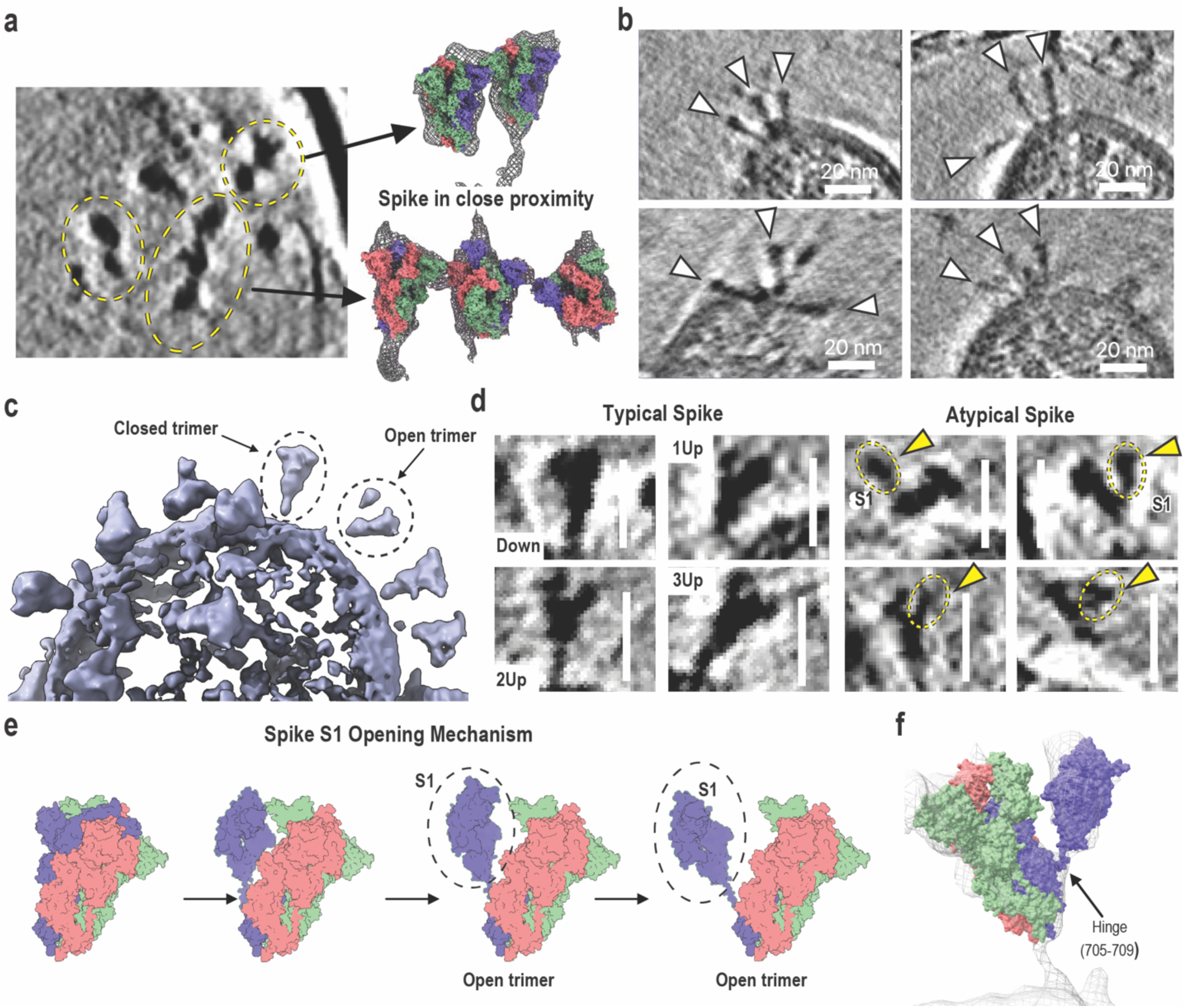
Distribution and atypical conformations of the spike protein. a) The 2D central slice of virus particle showing spikes in close proximity. Isosurface is also shown in mesh representation, wherein PDB 6ZGE ^10^ is fitted and shown in surface representation. b) Tomograms slices (7.5 nm thick) of four virions with clusters of postfusion spike structures. Postfusion conformations are highlighted with white arrowheads. c) The isosurface representation of several spike proteins on the virus surface, wherein the closed/packed and open trimers of prefusion states can be identified as shown by dashed lines. d) The magnified view of 2D central slice tomograms of the typical spike protein densities including Down, 1Up, 2Up and 3Up conformations, as well as atypical spike protein densities observed in active viruses. The detached S1 subunits are highlighted by dashed lines. e) Proposed schematic representation of the spike opening mechanism at prefusion states, in which one protomer S1 subunit undergoes a large conformational change. f) Simulation-based trimer model (surface representation) fitted to raw tomogram data of an open trimer (mesh representation). The length of scale bars in all images are 20 nm. **Figure 2–figure supplement 1.** Spike stem in close proximity and atypical conformations. **Figure 2–figure supplement 2.** Conformational flexibility in the spike protein isolated monomer. **Figure 2—video 1.** Zoomed view of atypical open-trimer spike protein from the reconstructed tomograms.

### A transient Open Trimer Spike Conformations

Both 247 prefusion and 44 postfusion conformations were observed across active virus particles. Figure 2c and d show all dominant forms of the spike protein observed in our dataset. This includes the prefusion Down, 1Up, 2Up and 3Up states, reflecting the inherent dynamical nature of the spike protein, in the absence of ACE2 or neutralizing antibody (nAb). Notably, besides typical conformations, a transient forms of the spike protein were also observed. Among these, spike protomers may undergo large conformational changes, forming transeint open trimer states. Fitting of S1 from monomer MD simulations to the detached density indicates that the observed motion involves the S1 subunit of single spike protomer. Accordingly, a range of atypical open trimer structures can be seen (Figure 2d, and 2e, Figure 2—video 1), likely exposing the S2 subunit. Note that in our analysis, any unidentified density on the viral surface, where spike proteins are expected, is most likely to represent a noncanonical or transient spike conformation. While the final count of 11 open trimers was based on visual assessment, we believe this number is likely an underestimate, as additional open trimers may be present but not readily identifiable due to orientation or resolution limitations. In fact, more open and identified types of trimer structures that may represent other transient states were also observed (Figure 2–figure supplement 1b).

Molecular Dynamics (MD) simulations of Spike monomer confirm the significant motion in the S1 domain, occurring at a new hinge region between residues 705 and 709 (Figure2f and Figure 2–figure supplement 2). This region includes a glycosylated Asn, which seems to promote and facilitate this hinge-like motion. Preliminary analysis of a larger dataset replicates the observation of a transient open trimers, confirming their formation. These results suggest a hidden motion of the spike structure, in which spike protomers are more flexible than observed in previous studies and can undergo large conformational opening. Consistently, previous HDX-MS studies using the recombinant ectodomain of the spike protein with a prefusion stabilization (2P mutations) suggested the presence of an alternative conformation, in which S2 residues are solvent exposed that can be targeted by nAbs, such as 3A3 and RAY53^24,25,28^. Likewise, high-speed AFM results pointed out the presence of large lateral motion in S1^27^. Hence, in active viruses, the typical closed/packed prefusion conformation is dominant but the spike protein itself has ability to spontaneously transit to atypical open-trimer conformations.

### Spike Average Conformation: Loosely Packed Structure

To gain further insights into the mechanism of the atypical-trimer formation, we performed subtomogram averaging (STA) analysis of the prefusion packed conformation, applying single-particle analysis of pseudosubtomograms as implemented in RELION 4.. 237 prefusion spike particles were manually picked (excluding open trimer and postfusion states). Around 28,000 tilt images (121 tilt per particle) were used for the 3D re-construction, with a voxel size of 1.884 Å. To avoid any bias, no previously published maps of the spike protein were used in the alignment and refinement processes. Figure 3a and Figure 3—video 1 show the sub-tomogram averaged map of the C3 symmetric spike protein, which was obtained with a global resolution of 14 Å (Figure 3–figure supplement 1, Supplementary file 1D, Figure 1). In addition, a C1 symmetry map was also obtained at a slightly lower global resolution of 17.0 Å, and this was essentially like the C3 symmetric structure but with less density in one protomer (Figure 3–figure supplement 1 a). The general features of the obtained maps are resembling those of previous cryo-EM studies. The overall map fits well with previous cryo-ET STA map (8.9 Å) that was obtained in the absence of ultracentrifugation^20^. A 2.6 Å resolution structure of the ancestral spike protein (PDB 6ZGE)^10^ fits adequately in the obtained low-resolution map (Figure 3a right). However, a better fit was obtained using the asymmetrically refined structure (PDB 7M0J) of the engineered spike (u1S2q) with quadruple mutations (A570L, T572I, F855Y, N856I)^34,35^ (Figure 3a left). The u1S2q spike show higher intrinsic mobility in comparison to 2P or 6P recombinant spike, which was evidenced by its high propensity of forming 1Up and 2Up conformations and sustained binding to the CR3022 antibody in similar fashion to highly exposed isolated RBD^35^. The fitting of the full spike structure (PDB: 6XR8)^6^ showed a better fit than 6ZGE, while the u1S2q mobile spike demonstrated the best fit.

**Figure 3.**
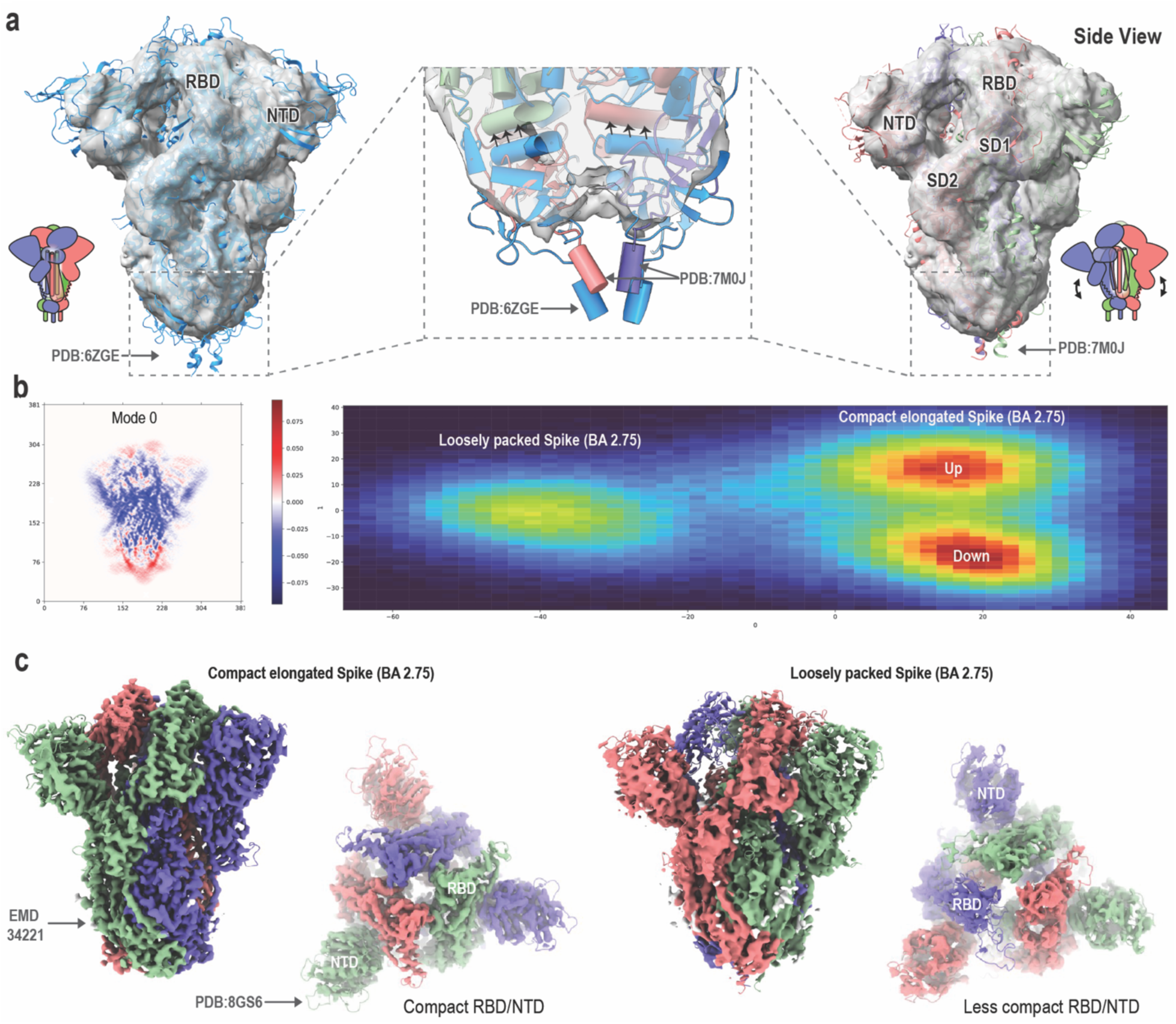
The spike protein averaged map obtained by STA analysis and cryo-EM. a) The sub-tomogram averaged map with the superimposition of cryo-EM structures of the spike trimer (PDB 7M0J; Right)) and (PDB 6ZGE; Left. Purple, green and pink colors were used to represent spike protomers. The zoomed-in view of the terminal part of spike protein averaged map (Bottom center) with fitted PDB structures, 7M0J (pink, green and purple ribbons) and 6ZGE (blue ribbon). Arrows in central panels indicate structural shifts from 6ZGE to 7M0J. b) Variability analysis of the BA 2.75 dataset showing minor conformation of the loosely packed trimers in agreement with STA map. c) Comparison of the previously published BA2.75 map (left) and model with the new loosely packed 3.72 Å map (right) and model (based on PDB:7M0J) of loosely packed spike conformation. **Figure 3–figure supplement 1.** STA map and spike distribution. **Figure 3–figure supplement 2.** Middle Helices and Charge distribution in Spike protomer. **Figure 3–figure supplement 3.** Vaccine design approaches. **Figure 3—video 1.** Sub-tomogram averaged structure fitted to engineered mobile spike protein.

In detail, the STA map shows two main differences from previous cryo-EM/cryo-ET maps. First, the overall map shows the formation of shorter and more bulged nature. Figure 3a highlights the fitting of 6ZGE and 7M0J to the proximal part of the stem helices, wherein the 7M0J structure shows an upward shift that better fit the averaged map. Previous principal component (PC) analysis of 165 protomers from 52 spike Down PDB structures, shows a similar behavior where PC1 and PC2 modes both show up and down motion in S2 with respect to S1, indicating structural variation within the available PDB structures^36^. Second, a missing density was observed at the S2 middle helices interface ( Figure 3–figure supplement 2a). The 7M0J structure, fitted to our cryo-ET map, features slightly bent helices. Charge analysis shows multiple regions with charged residues at the trimer interface (Figure 3–figure supplement 2b). A recent STA map of Env protein (9.1 Å resolution) in intact HIV particles also suggests flexibility in the middle helix region^14^. Notably, these structural features may allow for more flexible movement of the protomers and the formation of the above-mentioned open trimer structure as well as the transition to postfusion (Figure 4).

**Figure 4.**
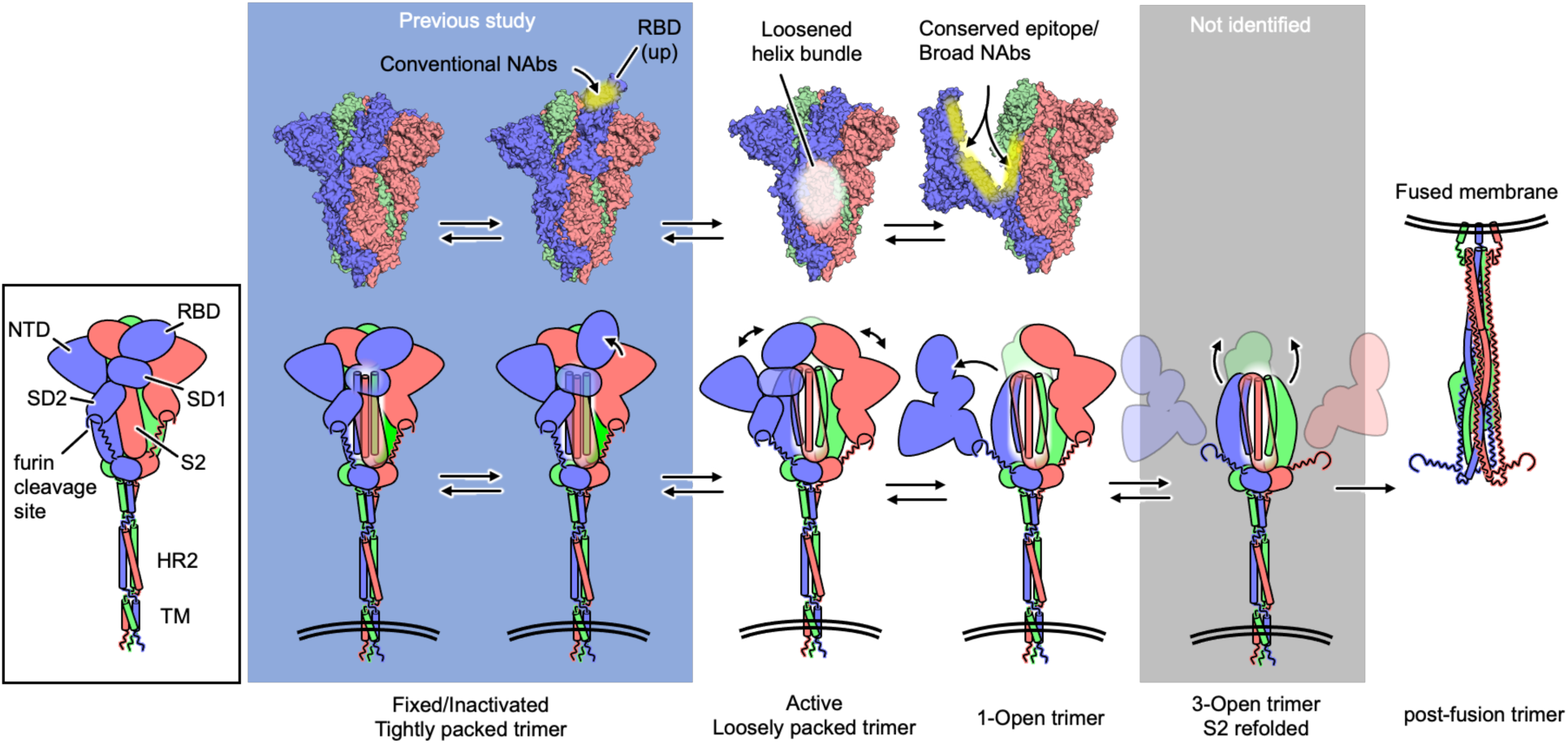
Proposed model of spike structural changes for the membrane fusion. (left panel) Schematic representation of the spike structure with the domain organization. The color of each protomer corresponds to Figure2e. (top left to center) New cryptic sites of the S2 subunit exposed in open-trimer conformations of the spike protein, potential targets for vaccine and antibodies/drugs. (bottom) Schematic presentation of the proposed fusion mechanism. Fixed or recombinant spikes protein take a closed or Up RBD structure (gray box). On the active virus, spike proteins exist as a loosely packed trimer, with a partially opened S1 subunit. When the ACE2 binding promotes the S2 subdomain exposure, the S2 subunit irreversibly refolds to the postfusion state to fuse the viral envelope with the host cell membrane.

### Loosely Packed Spike Conformation in Omicron BA.2.75

Considering the low resolution of the STA map, we re-analyzed our previously published cryo-EM data set of recombinant spike protein from Omicron BA.2.75^37^ to search for minor conformations of a loosely packed trimer. The Omicron dataset was selected because this variant, which carries multiple spike substitutions including the early D614G mutation, is known to exhibit enhanced structural flexibility and conformational heterogeneity. Figure 3b shows the results of continuous density variability analysis^38^, where mode 0 represents an upward motion of the spike conformation, transitioning from an elongated, tightly packed trimer to a shorter, loosely packed conformation. This conformation is highly distinct from typical Up and Down and accounts for less than 20% of the data set. Despite the proline mutation, loosely packed trimers occur as a minor population, highlighting the effect of stabilizing mutations on altering protein structure, as well as the limitations of traditional averaging schemes in cryo-EM, which may have overlooked such conformations.

Starting with the 7M0J model, the spike sequence was mutated and fitted to the obtained 3.72Å map (Figure 3–figure supplement 1c), showing the formation of more distant RBDs and NTDs in the S1 (Fig 3c). Notably, this conformation may allow for lateral motion of the S1, which is consistent with the observed lateral motion in active virions.

## Discussion

Mutations of the spike protein are a major underlying cause for immune escape and reduction of vaccine efficiency. The development of broadly neutralizing antibodies (bnAbs) is important for preparedness for future pandemics. The recent review of Zhou et al. summarized three major non-RBD sites targeted for bnAbs, (1) FP, (2) SD1 (the S1 subdomain 1) and (3) the C-terminal stem helix^39^. Our cryo-ET map shows that the spike protein is flexible enough for these sites to be to some extent exposed in open-trimer structures, possibly generating bnAbs that exhibit neutralizing activity but with less potency. Either cocktails or bispecific forms of bnAbs, which can recognize different epitopes of single spike protein spontaneously, may improve the activity and tolerate mutations^39^.

The development of long-term and broadly effective vaccines also requires new strategies to target highly conserved epitope regions^40,41^. Unlike RBD which is a hot spot for mutations, the S2 subunit shows higher conservation, wherein few mutations have emerged since the start of the pandemic. In fact, FP and heptad repeat regions (HR1 and HR2) of the S2 subunit are more conserved within all coronaviruses including SARS-CoV-1 and Middle East respiratory syndrome coronavirus (MERS)-CoV^41^. This emphasizes the potential of developing broadly effective vaccines that may cover current and future coronavirus outbreaks. However, in current vaccine development strategies, the S2 subunit is considered as inaccessible, wherein proline mutations are applied to further stabilize the prefusion state. On the contrary, our cryo-ET study here suggests the dynamic nature of the spike protein where parts of the S2 subunit are accessible (Figure 2d and Figure 4). As described above, bnAbs may exhibit less potent activity but vaccine-induced bnAbs would overcome this limitation, while the stabilization of the S2 subunit may be necessary. In fact, the development of a multivalent S2-based vaccine was recently shown to provide a wide effect against beta, delta and omicron variants as well as pangolin coronavirus^40^. Furthermore, a recent study has demonstrated that immunization with prefusion-stabilized S2 provide a broad neutralization effect^42^.

Besides the S2 subunit, the currently observed dynamics of the active SARS-CoV-2 virus (Figure 2–figure supplement 2) indicates the accessibility of the interface of S1 and S2 subunits. Note that these regions are conserved across SARS-CoV-2 variants. Indeed, the HDX-MS experiment of the recombinant spike protein also shows the increased accessibility of the two subdomains (SD1 and SD2) in the S1 subunit^25^. Collectively, accounting for spike dynamics and cryptic epitope formation may facilitate the development of broadly effective vaccines. For instance, a vaccine design based on the spike protomer may better account for spike protein dynamics, exposing cryptic epitopes (Figure 4 and Figure 3–figure supplement 3). MD simulations of the spike monomer of the Omicron BA.2 variant (Figure 2–figure supplement 2a and b) shows that a large range of motions in the S1 subunit can be observed in the absence of the trimeric complex, exposing cryptic pockets located at the interface of the S2 subunit, RBD, and the SD1 subdomain, which may induce a broader spectrum of immune effect. Indeed, monoclonal antibodies (mAbs) targeting cryptic epitopes, such as CR3022, P008-60, and sd1.040, can break up spike trimeric structure^43–45^. It is worth mentioning that an early study has reported the feasibility of using a monomeric spike protein of SARS-CoV-1 as a vaccine candidate^46^.

Unexpectedly, the active dynamics of the spike protein structure, especially the transeint open-trimer conformations, reveal benefits for fusion efficiency. This is in line with previously reported findings for hemagglutinin in influenza virus^47^ Some portion of open-trimer conformations likely expose the S2’ cleavage site for TMPRSS2 and cathepsins. In this sense, the spike protein possesses the inherent potential to allow the S1-S2 dissociation, inducing the exposure of the S2’ cleavage site. This process could be further facilitated by the ACE2 binding (Figure4). In retrospect, the effect of the D614G mutation would be anticipated to lead to a larger exposure. Such dynamic is expected to be altered by binding charged proteoglycans or changing local ionic strength. The spike protein in active virus is highly dynamic with multiple levels of flexibility including, membrane diffusibility, stalk region knee-like motion^18^, RBD Down to Up transition ^48^ and open-trimer formation^25^. In addition, it is highly susceptible to change in temperature, pH and even glycosylation^9,49^. Structural analysis using inactivated viruses or recombinant proteins is essential, however, the effect of mutations, chemical fixations, and trimerization tags shall be considered. Hence, it is crucial to study viral proteins in their native, active state. Cryo-ET at BSL3 facility is expected to be an indispensable method to tackle current and future pandemics.

## Materials and Methods

### Virus preparation

An ancestral SARS-CoV-2 strain, WK-521 (GISAID: EPI_ISL_408667), was kindly provided by Dr. Shimojima (National Institute of Infectious Diseases, Japan). The viral stock was prepared by the inoculation to Vero E6 cells constantly expressing human TMPRSS2 (Vero E6/TMPRSS2)^50^. At 2 days post-infection (dpi), the culturing supernatant was centrifuged at 440 × g for 2 min to remove the cell debris and stored at −80 °C as a virus stock. The titer of the stock was determined by plaque assay as previously described^51^. In brief, Vero E6/TMPRSS2 cells were inoculated with serial dilutions of virus stock for 1 h at 37 °C. The cells were overlaid with Dulbecco’s Modified Eagle Medium (D-MEM) containing 2% FBS and 0.5% Bacto Agar (Becton Dickinson). At 2 dpi, cells were fixed with 3.7% formaldehyde in phosphate-buffered saline (PBS) and stained with 1% crystal violet.

For the preparation of the cryo-ET sample, Vero E6/TMPRSS2 cells in a 6 well plate were inoculated with the virus at a multiplicity of infection (MOI) of 10 and incubated for 1 h at 37 °C. The media was then replaced with 400 μL of fresh Opti-MEM (Thermo Fisher Scientific). At 18 h post-infection, culture supernatant was collected and clarified with 0.45 μm filtration. The sample was kept at 37 °C and used immediately for grid preparation.

To prepare inactivated virus, paraformaldehyde was added to the native virus aliquot solution described above to a final concentration of 0.5% and incubated at 4 °C for at least 24 hours. As fiducial markers, 1 μL of 0.1% colloidal gold (10-nm diameter) suspension was placed behind the grid before loading the sample.

### Data collection

For cryo-ET, 2 μL of sample was applied on each side of a freshly glow-discharged QUANTIFOIL R 1.2/1.3 Cu 200-mesh grid (Quantifoil Micro Tools). The grid was blotted at once for 15 s from the back side and plunged into liquid ethane using EM GP2 (Leica Microsystems) at 18 °C and 90% humidity chamber setting. The micrograph images were collected on a Krios G4 (Thermo Fisher Scientific) operating at a voltage of 300 kV with an energy filter Selectris (slit width 10 eV) and a direct electron detector Falcon 4. Each micrograph was collected at a nominal magnification of 64,000× (pixel size of 1.88Å) with an exposure time of 0.73 s (dose of 1.5 e/Å^2^) at a underfocus of 0.2 μm using a Volta phase plate (VPP). A total dose of 150 e/Å^2^ was used. Tilt-series data collection was performed with a dose-symmetric scheme tilting by 1° steps from −60° to 60° using Tomography software (Thermo Fisher Scientific), Supplementary file 1C, Table 3. All experiments were approved by the local committees and Thermo Fisher Scientific in accordance with Japanese law. With regards to the inactivated virions, the grids were prepared with gold particles for better alignment and EM data were collected with same protocol with two magnifications of 64,000× (pixel size of 1.88Å) and of 81,000× (pixel size of 1.49Å), using 121° steps from −60° to 60°.

### Cryo-Electron Tomography Data Processing

42 micrographs were processed using the IMOD package^52^. Dose weighting was applied using alignframes funtion in Etomo. Patch tracking was used to align tilt images active virions, while Beadtrack was used for inactive ones. Ctfplotter was used to estimate defocus, phase shift, and astigmatism within the context of the contrast transfer function^53^. Prior to 3D construction, all images were CTF corrected, and dose filtered. Tomograms were binned 4x and reconstructed with weighted back projection with a voxel size of 7.536 Å. SIRT-like filtered tomograms were generated for visualization and particle picking. Reconstructed tomograms were also imported and filtered in EMAN2^54^. Virus particles were manually segmented in EMAN2 and used for isosurface visualization in Chimera-X software^55^. The two longest diameters of viruses and the thickness of ice layer were calculated using 3dmod.

291 spike proteins, from 46 virions, were manually picked from the 4-fold binned SIRT tomograms, including pre (247 particles) and postfusion (44 particles) states. The sub-tomogram averaging of the prefusion state was processed in Relion 4.0^29^, using 237 clearly visible prefusion particles (∼28000 tilt images). The reference-free alignment was performed, where the initial map was generated from the binned subtomograms and used subsequently for other refinements. After obtaining an adequately representative map, the C3 symmetry was invoked. A series of refinements were performed starting from 4x binning to 2x to unbinned. Finally, several cycles of the 3D refinement were performed at a voxel size of 1.884 Å on unbinned subtomograms. The CTF was refined using ctffind-4.1.14^56^ and 2D images alignment was improved in each refinement cycle. The cycles were repeated until no further improvement in resolution was observed. The final average map of the C3 symmetry was estimated to have 14.3 Å resolution at FSC = 0.143 cutoff in RELION4.0. To understand the effect of symmetries, a final cycle of refinement was performed with C1 symmetry, providing a 17.0 Å resolution map. The original RELION FSC curves including randomized phase is shown in Supplementary file 1D, Figure 1. The models of the spike protein, PDB 6ZGE and 7M0J, were rigid fitted in the C3 map using ChimeraX^55^. PyMOL software was used for visualizing the structures^57^.

### Cryo-Electron Microscopy Data Processing

The previously published cryo-EM dataset was re-analyzed using variability analysis in CryoSPARC 4.4.1^38,58^. A total of 3308 micrographs were used for analysis. A protocol similar to the previously published one was used^37^. Using Blob Picker 442, 33 particles with a box size of 382Å were extracted and used for processing, forming a 2.83Å map of the typical down conformation. To successfully observe minor conformations, clear dominant down conformations were excluded (183,884 particles) from the dataset, and variability analysis was performed using 290,853 particles. Ten clusters were constructed, and a map was refined for each. The refined maps were used as input for 3D classifications to better redistribute particles using all particles, including the previously excluded ones (427,000 particles). Subsequently, new variability analysis was performed using a mask from the previous 3D class step. Using mode 0, the dataset was further clustered into 3 clusters to separate loosely packed trimers from the typical down and up conformations, resulting in 119,653 particles. Those particles were further classified and refined using 2D class, heterogeneous refinement, and variability analysis. Finally, an FSC 3.72Å resolution map was obtained using 29,413 particle.

### MD simulations of Spike monomer

The monomeric spike protein model was based on the Omicron BA.2 Cryo-EM structure (PDB 7UB0)^59^. The starting PDB structure has the following missing regions (70-80, 146-162, 178-186, 246-261,623-640 and 678-688), the galaxyFill in CHARMM-GUI was used to model missing regions. CHARMM-GUI was also used to add ions, water molecules for preparing final simulation model^60^. Glycan and disulfide bonds were added in a similar fashion to previous study^48^. Considering the potential large motion in monomer spike protein, a large box size was used with total number of atoms of 742586, including spike protomer (resid 17-1139), 19 glycans, 683 Na and 685 Cl ions in a water box with side length of 195 Å. All simulations were performed using GENESIS 2.0 Beta MD software on Fugaku supercomputer^61,62^. Three independent simulations for 100 ns each was performed using CHARMM 36M force field for protein residues and CHARMM TIP3P for ions and water molecules^63,64^. VMD software was used for trajectory visualization^65^.

System equilibrations were performed in a step manner, where we start with restraint on backbone and glycan dihedral and gradually remove the restraints. Similarly, the system was well equilibrated in NVT and NPT. First, minimization with 10,000 steps is performed with restraints on backbone atoms and glycan dihedrals. This is followed by heating in the NVT ensemble with the VVER integrator for 52,000 steps at a 1 fs time step, keeping the same restraints. Next, there are two equilibration phases: the first in NVT for 250,000 steps with a 2 fs time step and the second in NPT for 250,000 steps, where restraints are removed. Finally, two additional equilibration steps in NPT and NVT are done with 2.5 fs and 3.5 fs time steps respectively, without any restraints. Starting from equilibration 4, three independent simulations were performed for around 100 ns for run1 and run3. Due to the large observed RMSD in Run2, we further elongated the simulation for 200 ns. The 200 ns show same behavior, so only the first 100 ns was added to results sections for consistency with the other two runs. RMSD of all Cα atoms was performed upon fitting S2 Cα atoms only (residues 854-1135).

## Acknowledgments

We thank Hiroshi Kida, Tatsuya Zenko, Atsushi Furukawa, Takashi Tadokoro, Robert Parsons, Peijun Zhang, Fabian Eisenstein, Radostin Danev, Makoto Takeda for their support and discussions. We also thank members of Facilities Division of Hokkaido University and Thermofisher Scientific (Yoshihiro Tomatsu, Kazuhiro Aoyama, Yu Yong) for their technical assistance. This research used computational resources of the supercomputer Fugaku provided by the RIKEN Center for Computational. The computer resources were provided to H.D (Project ID: hp220087, hp230052, hp240149, hp250074).

## Funding

Japan Agency for Medical Research and Development (AMED) [JP17am0101093 (K.M.)]

Japan Agency for Medical Research and Development (AMED) [JP20ae0101047 (K.M.)]

Japan Agency for Medical Research and Development (AMED) [JP21fk0108463 (K.M.)]

Japan Agency for Medical Research and Development (AMED) [JP22ama121037 (K.M.)]

Japan Agency for Medical Research and Development (AMED) [JP223fa627005 (K.M, K.T, M.S, Y.O, and H.S.)]

Japan Agency for Medical Research and Development (AMED) [JP23ama121001 (T.S.)]

Hokkaido University Biosurface project (K.M.)

Japan Science and Technology Agency (JST) CREST grant number JP22gm1810004 (K.M), JPMJCR20H8 (H.F.)

Japan Society for the Promotion of Science (JSPS) Strategic Young Researcher Overseas Visits Program for Accelerating Brain Circulation (K.M.)

The Scientific Research on Innovative Areas and International Group from the MEXT/JSPS KAKENHI [JP20H05873 (K.M)]

Ministry of Health, Labour and Welfare (MHLW) [23HA2010 (H.S.)]

Takeda Science Foundation (K.M.)

Hokkaido University DX Doctoral Fellowship (Y.A.)

Hokkaido University EXEX Doctoral Fellowship (T.N., and T.S.)

## Author contributions

Designed the study: H.F., H.M.D., and K.M

Set up BSL3 Cryo-EM facility: H.F., S.K., Y.S., H.S. and K.M.

Assisted to set up and consult about BSL3 Cryo-EM facility: J.T.H. and D.I.S.

Prepared virus solution: H.F., K.T, M.S., Y.O., and H.S

Performed cryo-ET data collection: H.F., S.K., K.T., Y.A., and H.S.

Performed cryo-ET analysis: H.M.D., T.N., and T.S.

Advised Cryo-ET analysis S.K., J.T.H., and K.M.

Designed and prepared unique device of sample transfer for Cryo-ET: A.S. T.S., and K.M.

Wrote the manuscript: H.M.D., H.F., and K.M.

## Resources Availability

Various constructed tomograms have been deposited in the Electron Microscopy Data Bank (EMDB) under the following accession code EMD-39421. Likewise, the C3 symmetry STA map has been deposited in the EMDB (EMD-39422). MD simulation models (PDB and PSF) of the spike protein in its monomeric form have been deposited in Zenodo (https://doi.org/10.5281/zenodo.13968676).

## Appendix

### Key resources table

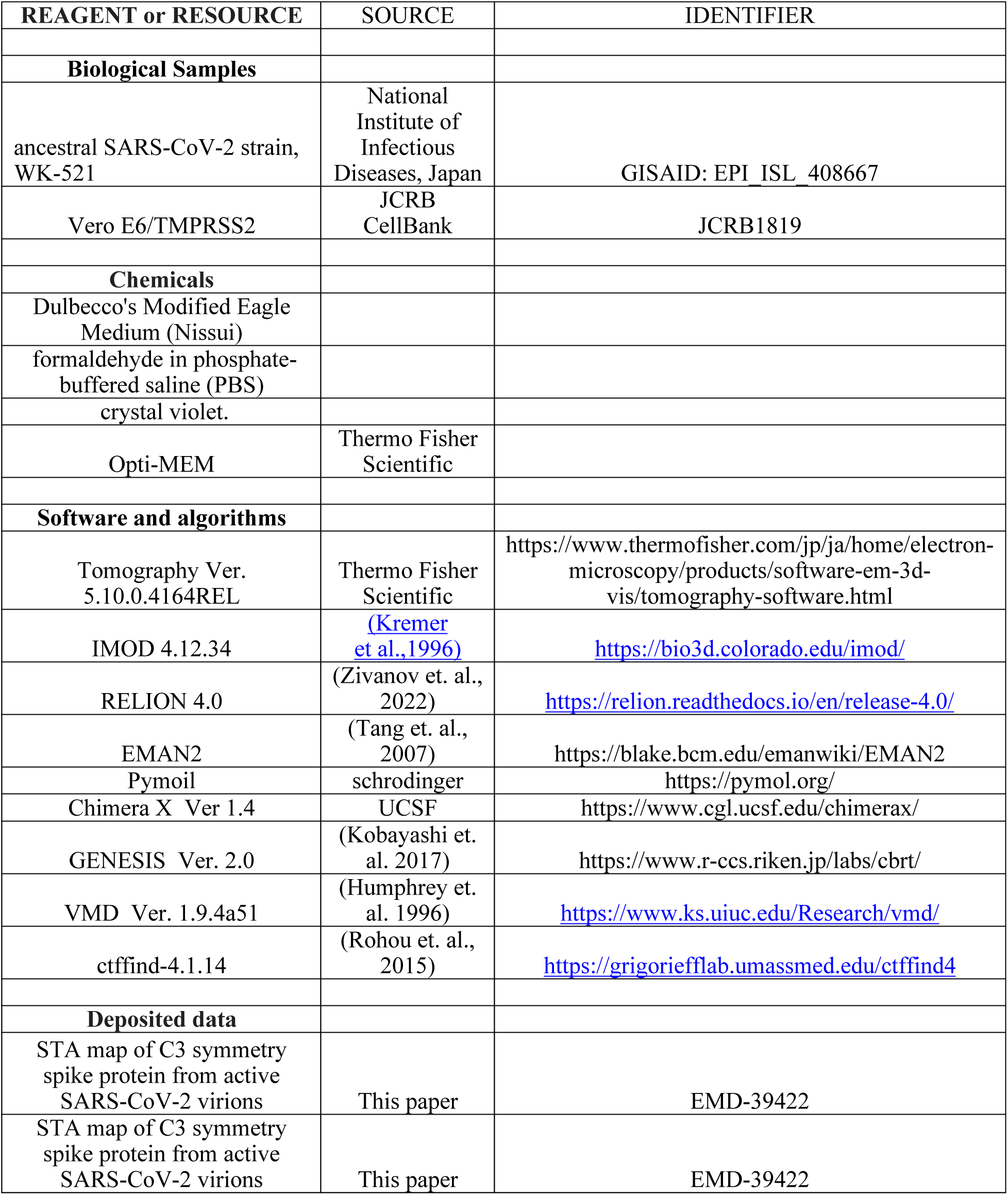

| REAGENT or RESOURCE | SOURCE | IDENTIFIER |
| --- | --- | --- |
| **Biological Samples** |  |  |
| ancestral SARS-CoV-2 strain, WK-521 | National Institute of Infectious Diseases, Japan | GISAID: EPI_ISL_408667 |
| Vero E6/TMPRSS2 | JCRB CellBank | JCRB1819 |
| **Chemicals** |  |  |
| Dulbecco's Modified Eagle Medium (Nissui) |  |  |
| formaldehyde in phosphate-buffered saline (PBS) |  |  |
| crystal violet. |  |  |
| Opti-MEM | Thermo Fisher Scientific |  |
| **Software and algorithms** |  |  |
| Tomography Ver. 5.10.0.4164REL | Thermo Fisher Scientific | https://www.thermofisher.com/jp/ja/home/electron-microscopy/products/software-em-3d-vis/tomography-software.html |
| IMOD 4.12.34 | (Kremer et al., 1996) | https://bio3d.colorado.edu/imod/ |
| RELION 4.0 | (Zivanov et. al., 2022) | https://relion.readthedocs.io/en/release-4.0/ |
| EMAN2 | (Tang et. al., 2007) | https://blake.bcm.edu/emanwiki/EMAN2 |
| Pymol | schrodinger | https://pymol.org/ |
| Chimera X Ver 1.4 | UCSF | https://www.cgl.ucsf.edu/chimerax/ |
| GENESIS Ver. 2.0 | (Kobayashi et. al. 2017) | https://www.r-ccs.riken.jp/labs/cbrt/ |
| VMD Ver. 1.9.4a51 | (Humphrey et. al. 1996) | https://www.ks.uiuc.edu/Research/vmd/ |
| ctffind-4.1.14 | (Rohou et. al., 2015) | https://grigoriefflab.umassmed.edu/ctffind4 |
| **Deposited data** |  |  |
| STA map of C3 symmetry spike protein from active SARS-CoV-2 virions | This paper | EMD-39422 |
| STA map of C3 symmetry spike protein from active SARS-CoV-2 virions | This paper | EMD-39422 |

