## Supplemental Figures for "Native spike flexibility revealed by BSL3 Cryo-ET of active SARS-CoV-2 virions"

**This file includes:**
Figure supplement (7 figures and 5 videos)

### Supplemental Figures

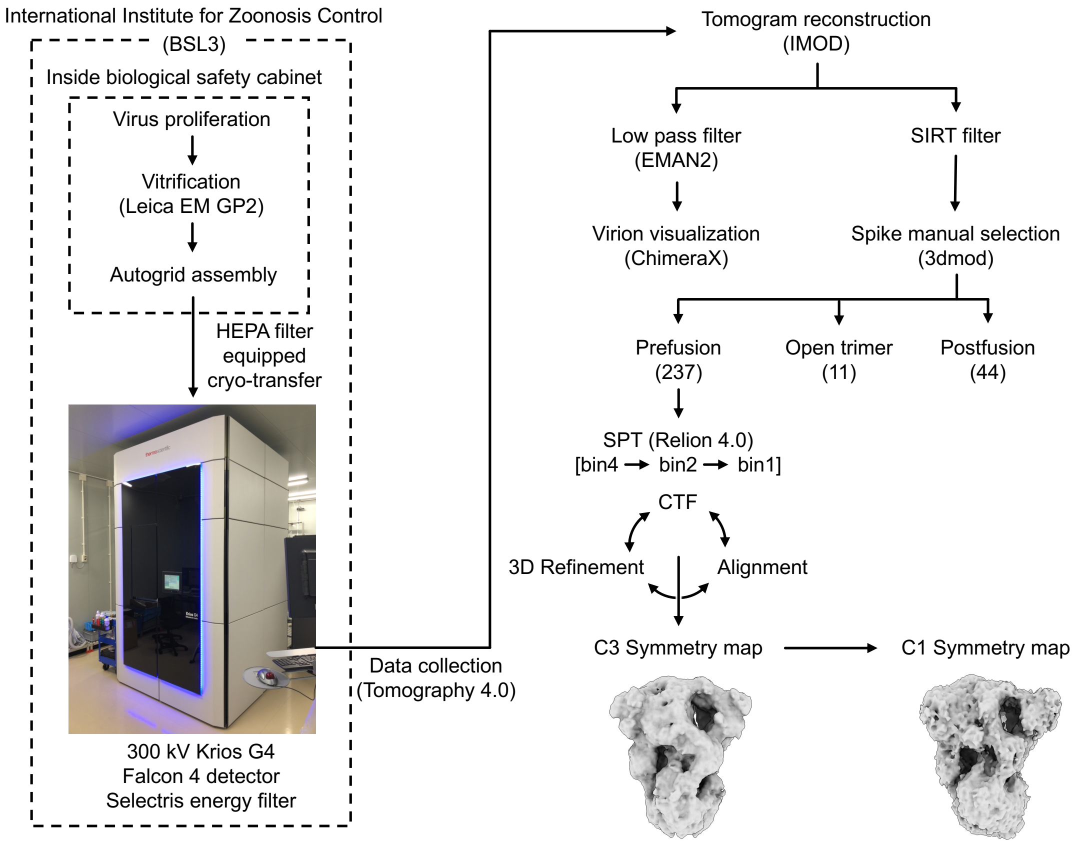


**Figure 1–figure supplement 1. BSL3 Cryo-EM facility at Hokkaido University and Cryo-ET analysis scheme.** Krios G4 was introduced in the BSL-3 room which is constantly controlled negative pressure and temperature with air circulation through HEPA filter. Sample vitrification using EM GP2 (Leica) and autogrid assembly were all performed in the safety cabinet. Autogrids were then transferred to the Krios G4 using a proprietary developed cryotransfer system equipped with a HEPA filter. After the measurement, the grid was taken out without thawing under the same procedure, immersed in 70% ethanol, and then autoclaved. For maintenance, the entire enclosure of the Krios G4 vacuum system was heated to 60$^{o}$C for over three hours using an installed heating system, and the room interior was sterilized by hydrogen peroxide vapor. The room sterilization was confirmed by biological indicator killing. Data were collected using Tomography 4.0. Subsequently, Tomograms were constructed using IMOD suite of programs. Spike proteins were manually selected, and STA was performed with Relion 4.0 software. Initially C3 symmetry map was obtained and subsequently and C1 map was constructed.

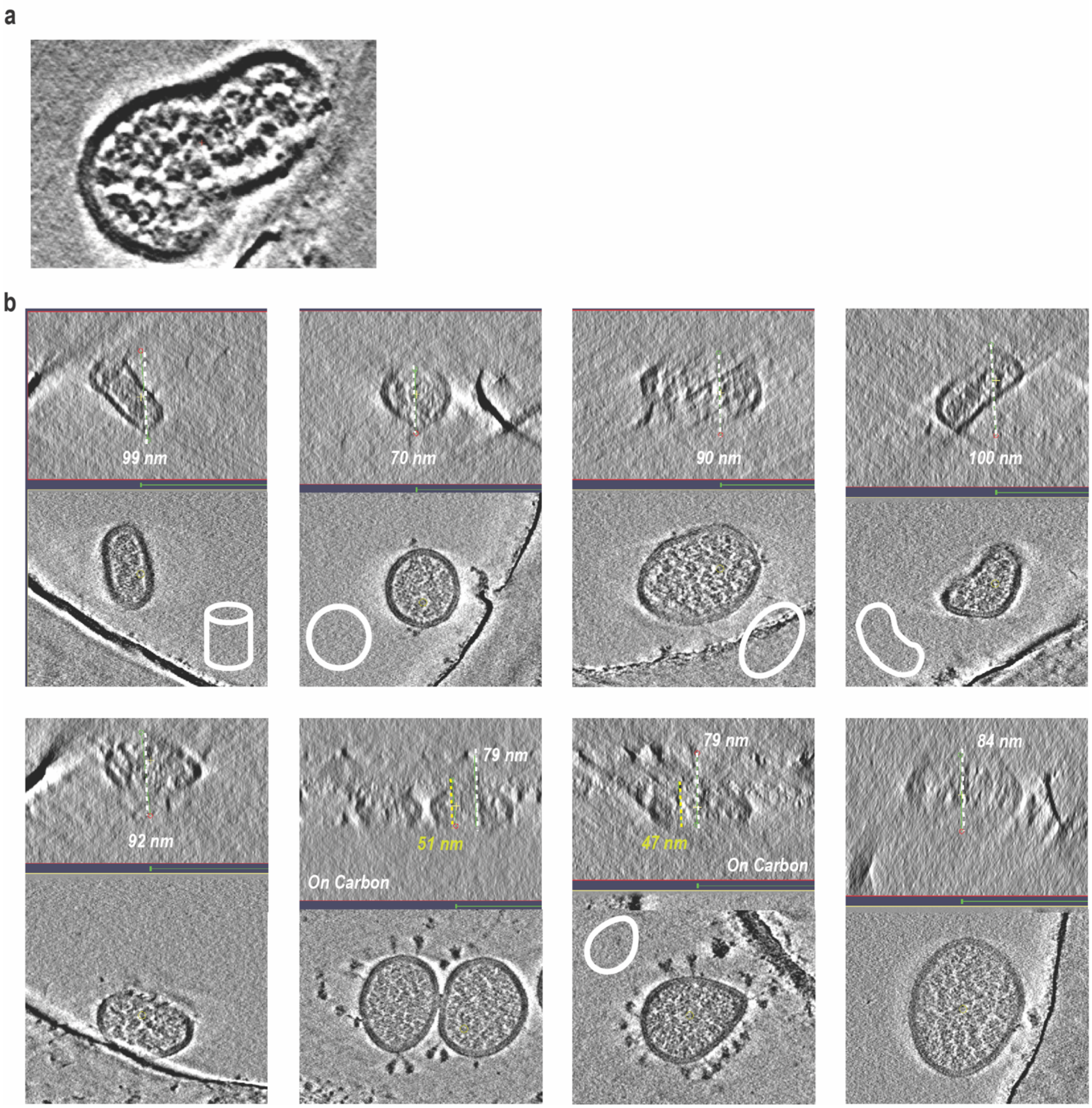


**Figure 1–figure supplement 2. Particle morphology and ice thickness.** a)Tomogram slice of two virion particles undergoing fusion process. b) Side and central sections through the cryo-ET tomogram of the ancestral virions' particles with different morphology. Ice thickness is highlighted by dotted line and reported in nm. Yellow and white lines represent virus and ice thickness, respectively.

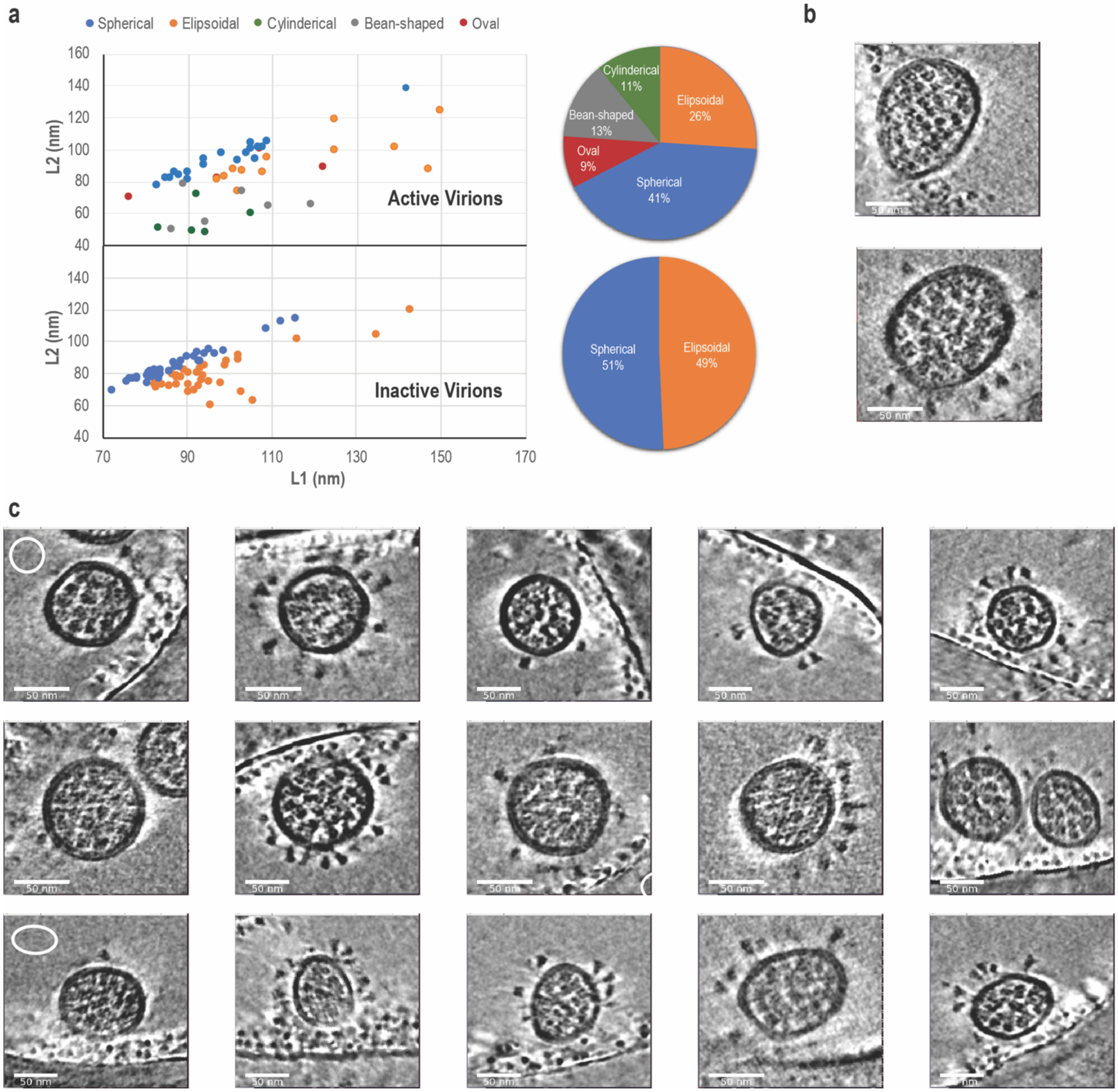


**Figure 1–figure supplement 3. Morphology of inactive virions.** a) Comparison of measured diameters between active (Top) and inactive (Bottom) virions, b) Central slice of the two largest virions in the inactive dataset, c) Central slices of inactive virions, demonstrating similarity with previous studies of fixed virions. The length of scale bars in all images are 50 nm.

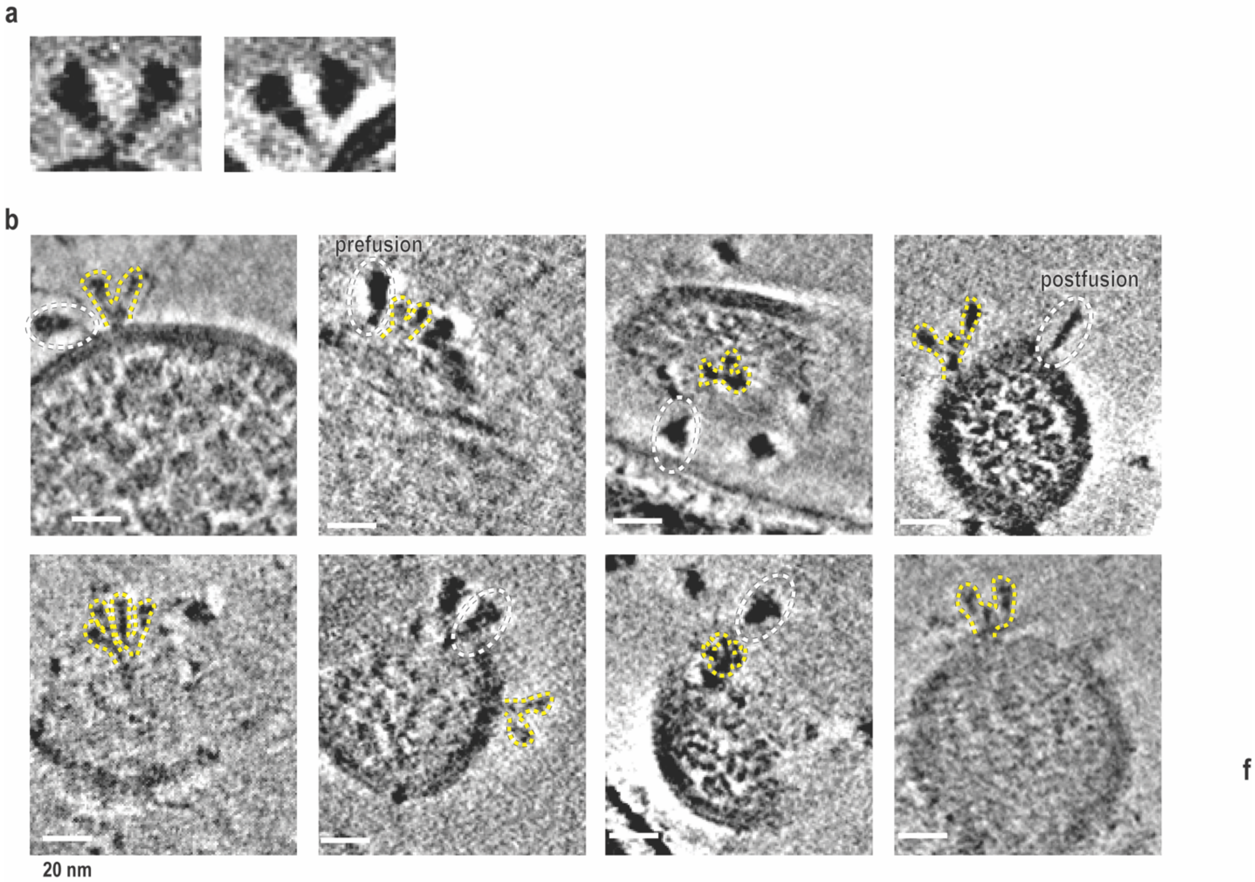


**Figure 2–figure supplement 1. Spike stem in close proximity and atypical conformations.** a) 2D slice of full spike protein structures with stems in close proximity. b) Zoomed out views of 2D slices of distinct unidentified atypical spike conformations that may exhibit monomeric or fully open-trimer conformations. Atypical spikes are highlighted by yellow dashed lines. For comparison, nearby typical prefusion and postfusion spikes are highlighted by while dashed lines. Scale bar of 20 nm is also shown.

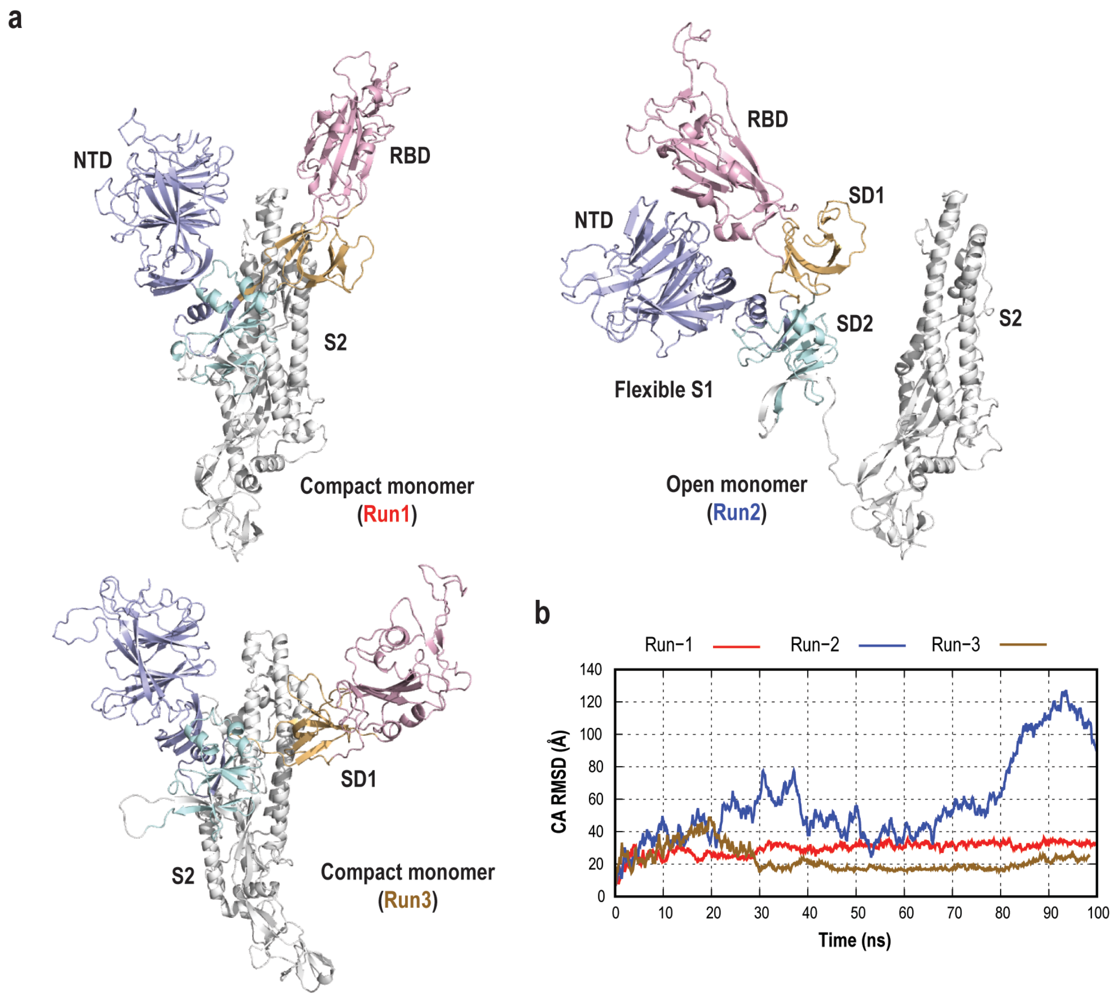


**Figure 2–figure supplement 2. Conformational flexibility in the spike protein isolated monomer.** a) Cartoon representation of different conformations of the head region of spike protomer obtained from three independent MD simulations. Two simulations form compact monomers conformations, while one simulation shows the opening of the S1 subunit (open monomer). b) Root mean square deviations (RMSD) of the C$\alpha$ atoms upon fitting S2 region. All simulations show large deviation from a starting structure due to the motion of the S1 subunit.

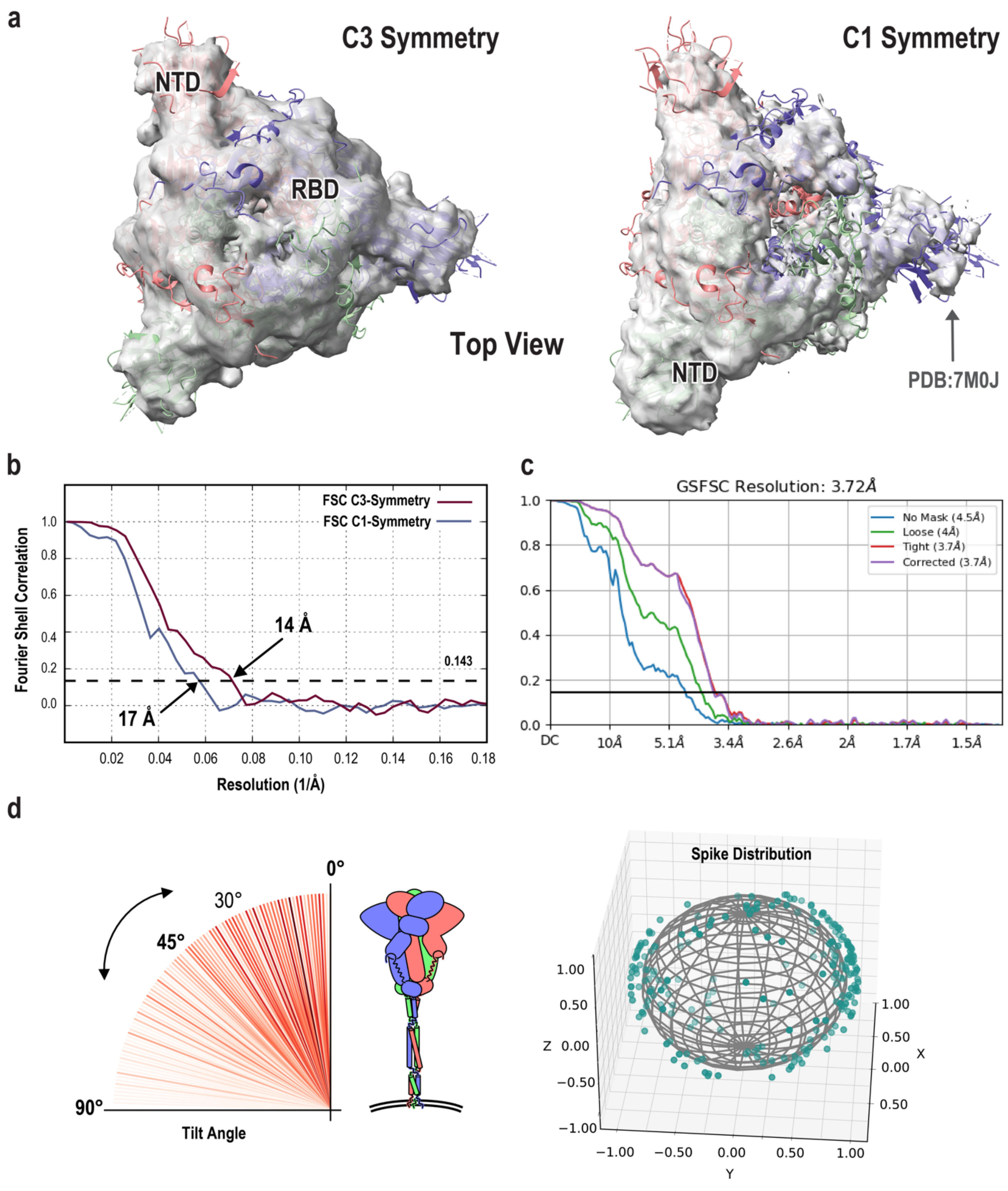


**Figure 3–figure supplement 1. STA map and spike distribution.** a) Top views of the sub-tomogram averaged maps of the C3 symmetry (top) and the C1 symmetry (bottom) fitted with prefusion-state spike protein (PDB 7M0J). b) Gold-standard Fourier shell correlation (FSC) curves for C3 and C1 symmetrized maps. Dotted line indicates the 0.143 FSC cutoff. c) FSC curves for cryo-EM date of loosely packed trimer in BA. 2.75. d) Left: Spike protein tilt angles distribution, showing an average tilt angle of 30°. A Cartoon representation of spike protein is also shown.

Right: prefusion state orientation distribution map with normalized distances, shown as green dots. A sphere at fixed distance of 0.9 is also shown as grey contour.

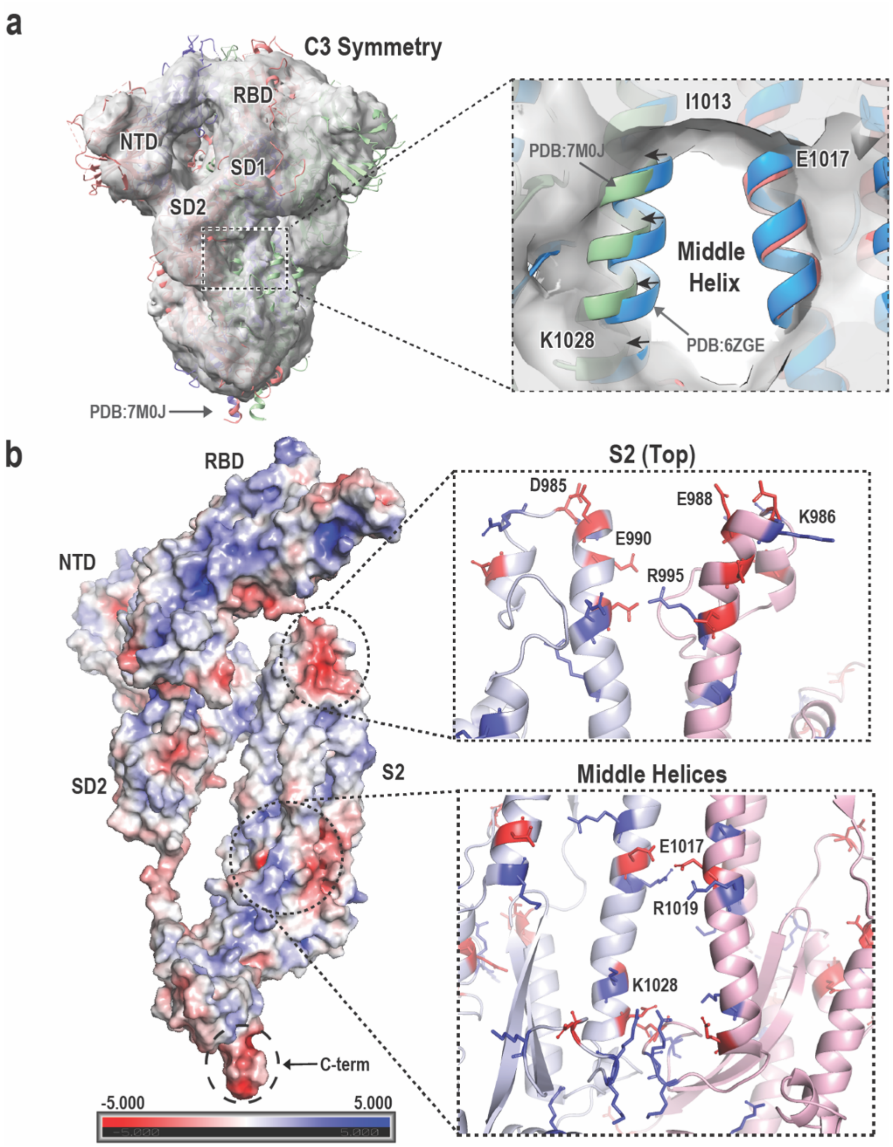


**Figure 3–figure supplement 2. Middle Helices and Charge distribution in Spike protomer.** a) STA map with fitted PDB structure, highlighting fitting of two different PDBs to the middle helix region. b) Electrostatic surface representation of the spike protein protomer from the high-resolution structure (PDB 6ZGE). Highly charged regions are highlighted with dotted lines. Ribbon representation of the middle and top regions of S2 interface from two protomers with highlighted charged residues. Blue and red sticks represent positively and negatively charged amino acids, respectively.

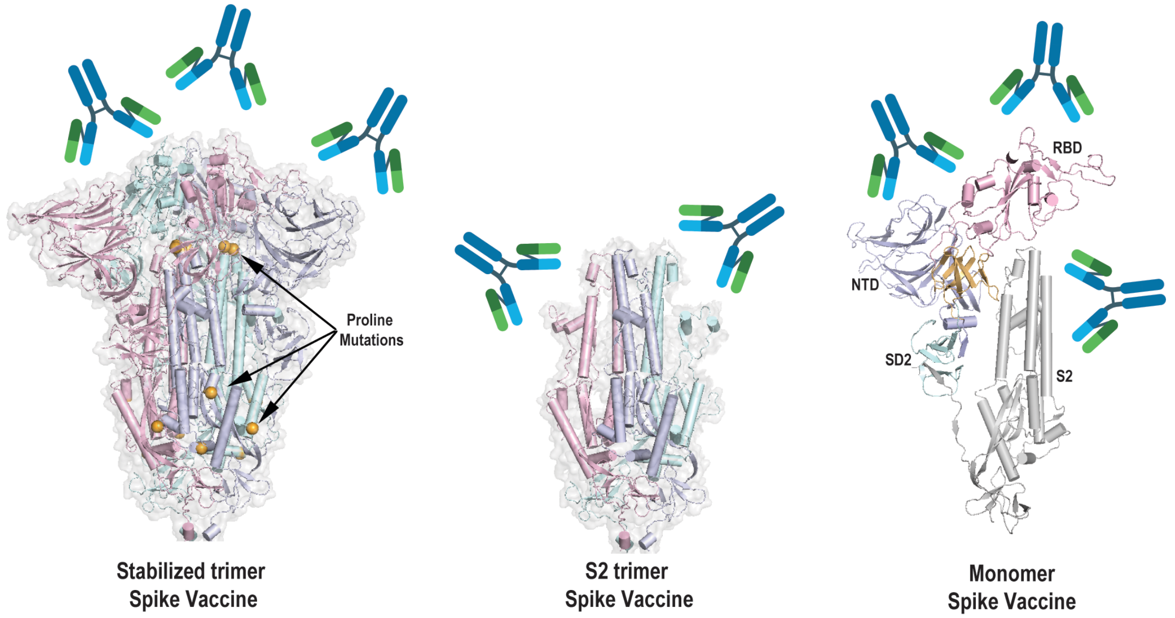


**Figure 3–figure supplement 3. Vaccine design approaches.** Left: The cartoon representation of second generation of SARS-CoV-2 Vaccines that include 6 proline mutations (shown as orange spheres), inducing mainly RBD and NTD neutralizing Abs. Middle: S2-based vaccine approach that allow for targeting the S2 region. Right: Monomer-based vaccine approach to account for protein dynamic and target both S1 and S2 regions. The cocktail approach of S1- and S2-based

**Supplemental Movies**

Figure 1—video 1.
2D and 3D Reconstruct of several tomograms of active SARS-CoV-2 virions.

Figure 1—video 2.
Large variations in virion sizes.

Figure 1—video 3.
Two virion particles undergoing fusion.

Figure 2—video 1.
Zoomed view of atypical open-trimer spike protein from the reconstructed tomograms.

Figure 3—video 1.
Sub-tomogram averaged structure fitted to engineered mobile spike protein.
